# The sequence of the extruded non-template strand determines the architecture of R-loops

**DOI:** 10.1101/576561

**Authors:** Yeraldinne Carrasco-Salas, Amélie Malapert, Shaheen Sulthana, Bastien Molcrette, Léa Chazot-Franguiadakis, Pascal Bernard, Frédéric Chédin, Cendrine Faivre-Moskalenko, Vincent Vanoosthuyse

**Author notes:** To whom correspondence should be addressed: Vincent Vanoosthuyse. Correspondence may also be addressed to Cendrine Faivre-Moskalenko.

## Abstract

Three-stranded R-loop structures have been associated with genomic instability phenotypes. What underlies their wide-ranging effects on genome stability remains poorly understood. Here we combined biochemical and atomic force microscopy approaches with single molecule R-loop footprinting to demonstrate that R-loops formed at the model *Airn* locus *in vitro* adopt a defined set of three-dimensional conformations characterized by distinct shapes and volumes, which we call R-loop objects. Interestingly, we show that these R-loop objects impose specific physical constraints on the DNA, as revealed by the presence of stereotypical angles in the surrounding DNA. Biochemical probing and mutagenesis experiments revealed that the formation of R-loop objects at *Airn* is dictated by the sequence of the extruded non-template strand, suggesting that R-loops possess intrinsic sequence-driven properties. Consistent with this, we show that R-loops formed at the fission yeast gene *sum3* do not form detectable R-loop objects. Our results reveal that R-loops differ by their architectures and that the organization of the non-template strand is a fundamental characteristic of R-loops, which could explain that only a subset of R-loops is associated with replication-dependent DNA breaks.

## INTRODUCTION

DNA:RNA hybrids have emerged as important regulators of genome stability and gene expression (reviewed in (1)). One possible source of DNA:RNA hybrids in the genome is the formation of co-transcriptional R-loops, where the nascent RNA hybridizes with its DNA template through Watson-Crick interactions, whilst the non-template DNA strand remains single-stranded (reviewed in (1)). A number of DNA/RNA helicases as well as the two ribonucleases H (RNase H1 & H2) target DNA:RNA hybrids and thereby maintain R-loop formation to low levels (reviewed in (1)). However, when these activities fail, wide-ranging evidence suggests that the resulting stabilization of R-loops interferes with the stability of replication forks during DNA replication and is associated with DNA damage and genome instability (2–5). The molecular mechanisms leading to DNA damage are still poorly understood but a recent study proposed that only a small subset of R-loop-forming regions (~10%) actually induces DNA breaks after DNA replication (2). To explain this observation, it was proposed that the chromosomal context in which R-loops form must contribute to the accumulation of DNA damage. An alternative and equally possible explanation could be that there are in fact different types of R-loops with different intrinsic properties, with some imposing greater physical constraints on the surrounding DNA than others. Here we test this possibility using Atomic Force Microscopy (AFM) to monitor the impact of R-loop formation on the surrounding DNA.

Both *in vitro* and *in vivo* observations show that the propensity of a transcription unit to form R-loops is determined at least in part by the sequence of the non-template strand. In particular, G-skewed non-template strands containing stretches of consecutive G residues significantly increase the probability of forming co-transcriptional R-loops (6–8). Although strong determinants of R-loop formation have been identified, little is known about the actual architecture of R-loops. In particular, how the extruded single-stranded DNA (ssDNA) is organized and stabilized has not yet been thoroughly addressed experimentally. It is conceivable that it could recruit specific proteins and/or form secondary structures, opening up the possibility that R-loop architecture is variable and dependent on several parameters such as size and/or sequence. For instance, it was reported previously that R-loop formation at the murine Sγ3 immunoglobulin switch regions allowed the extruded non-template strand to assemble G-quadruplex structures (9, 10), although this is controversial as others failed to detect G-quadruplex formation in the same conditions (11, 12). To date, an in-depth assessment of R-loop architecture and its potential diversity is missing.

Here we use a combination of single molecule approaches to study the architecture of transcription-dependent R-loops formed *in vitro*. We demonstrate that the extruded non-template single-stranded DNA (ssDNA) strand determines the architecture of R-loops and their physical properties and that R-loop architecture can be manipulated using site-directed mutagenesis. This opens the possibility that the extruded ssDNA is a distinguishing feature of R-loops *in vivo*.

## MATERIAL AND METHODS

### Plasmid preparation

The plasmids used in this study are shown on Figure S1. *Airn* and the *Airn G-stretch* mutant were synthesized (GeneCust) and cloned into pUC57. *sum3* was amplified from genomic DNA using the forward primer Xba_sum3 FW that contains the sequence of the T3 promoter (5’-GCGTCTAGAATTAACCCTCACTAAAGGGAATGAGCGACAATGTACAGC-3’) and the reverse primer Bam_sum3 RV (5’-GCGGGATCCTTACCACCAGGATTGAGCAC-3’) and cloned into the pUC57 plasmid using standard protocols. pDR18 (11) containing 4 repetitions of the immunoglobulin Sγ3 switch regions was a kind gift from Dr Michael R. Lieber (University of Southern California). The plasmids were purified using the NucleoBond^®^ XtraMidi kit (Macherey-Nagel) according to the manufacturer’s recommendations.

### *In vitro* transcription of R-loops

1875 ng of circular plasmids were incubated at 37°C with 50 U of T3 RNA Polymerase (Promega) in a transcription buffer (40 mM Tris-HCl pH 7.9, 6 mM MgCl_2_, 2 mM spermidine, 10 mM NaCl, 20 mM DTT, 0,05% Tween-20) containing 0,25 mM of rATP, rCTP, rUTP and rGTP (Promega). After 30 minutes at 37°C, T3 was inactivated by incubating the reaction at 65°C for 10 minutes. When specified, the NaCl in the reaction buffer was replaced by 40 mM KCl or 40 mM LiCl. 750 ng of transcribed plasmids were then incubated in the CutSmart Buffer (New England Biolabs) containing appropriate restriction enzymes and 2 U of RNase H. pFC53 was digested with ApaLI, pUC57-Airn was digested with Kpnl and Hindlll and pUC57-sum3 was digested with BamHI and Xbal. After 140 minutes at 37°C, the NaCl concentration was brought to 500 mM and 0,4 μg of RNase A was added for 20 minutes at 37°C to digest soluble RNAs. The DNA was then purified using chloroform and isopropanol precipitation and resuspended in 10 mM Tris. The DNA was run on 0,8% agarose gels prepared in TBE 1X without intercalating agent.

### Nuclease P1 treatment

After *in vitro* transcription and restriction digest, the NaCl concentration was brought to 500 mM and 10 U of nuclease P1 (reference N8630, SlGMA) was added. The mixture was incubated at 50°C for 45 minutes. The DNA was subsequently purified using choloroform extraction and isopropanol precipitation.

### Dot blot analysis

Serial dilutions of DNA (80 ng, 40 ng, 20 ng, 10 ng, 5 ng) were spotted on Hybond^TM^-N+ membrane (Amersham RPN203B) and cross-linked twice with UV (0,12J). To quantify DNA:RNA hybrid formation, the membrane was first blocked in MES/BSA buffer (100 mM MES pH 6,6, NaCl 500 mM, 0,05% Triton, 2 mg/mL BSA) for 30’ at RT and then incubated overnight in 0,02 μg/mL of S9.6 antibody. After extensive washes in MES/BSA buffer, exposure to a secondary antibody and ECL revelation was performed using SuperSignal West Femto Maximum Sensitivity Substrate (ThermoScientific 34096) according to standard protocols. To quantify dsDNA, the membrane was first blocked in PBS 1X 0,02% Tween-20 containing 5% Milk and incubated overnight with the dsDNA-specific antibody (ab27156 Abcam, dilution 1/1000). To quantify G4 formation, the membrane was first blocked in BG4 buffer (13) (25 mM Hepes, 10 mM NaCl, 110 mM KCl, 130 mM CaCl_2_, 1 mM MgCl_2_) containing 1% BSA and incubated overnight with the G4-specific antibody (BG4, MABE917, Merck) at 500 ng/mL in BG4 buffer containing 1% BSA. After extensive washings in PBS 1X 0,02% Tween-20, the membrane was incubated for 1 hour with an anti-Flag antibody (M2, Sigma, dilution 1/1000) at room temperature, followed by exposure to a secondary antibody and ECL revelation using SuperSignal West Femto Maximum Sensitivity Substrate (ThermoScientific 34096). As a positive control, a ssDNA oligonucleotide corresponding to a G4-forming sequence of the *cmyc* promoter (5’-GAGGGTGGGGAGGGTGGGGAAGG-3’) was incubated at 94°C for 5’ in either 10 mM Tris pH 7.4, 100 mM KCl (KCl G4 buffer) or in 10 mM Tris pH 7.4 100 mM LiCl (LiCl G4 buffer) and then let to cool down at room temperature for 1 hour. 30 ng of DNA (oligonucleotide or plasmid) was then spotted on the membrane.

### Atomic Force Microscopy

5 ng of DNA were diluted in TM buffer (10 mM Tris pH 7.4, 5 mM MgCl_2_) and loaded on mica disks previously cleaved using Scotch tape to obtain a flat surface. After 2 minutes, the sample was washed with 1 mL of ultrapure water and dried gently using a nitrogen flow. The samples were imaged using an Atomic Force Microscope (Bruker, Nanoscope V Multimode 8 model) using the Tapping Mode in air. Typical AFM of 3μm*3μm (512*512 pixels^2^) were acquired at a rate of 2 Hz using TESP or DLCS cantilever with a resonant frequency around 300 kHz. All images were edited using NanoScope Analysis program to remove the tilt and low noise frequency in the image, before further analysis (see Supporting material).

### SMRF-seq based R-loop footprinting

After *in vitro* transcription, the DNA was treated with sodium bisulfite under non-denaturing conditions (no denaturation and incubation at 37°C for 3 hrs) using the Zymo Lightning bisulfite modification kit (Zymo Research). Following sample clean-up, the DNA was amplified using native primers flanking the R-loop prone region and Pacific Biosciences sequencing libraries were built directly from PCR amplicons. Upon library validation, sequencing was performed on pooled barcoded libraries on a Pacific Biosciences RSII instrument. Following sequencing, the Gargamel computational pipeline was used to identify SMRF footprints by tracking significant patches of C to T conversion across high quality circular consensus single reads as described in (14).

## RESULTS

### R-loop formation at *Airn* allows the folding of the non-template strand into secondary structures

The non-template strand of the mouse *Airn* non-coding RNA (ncRNA) was previously shown to display a strong G-skew over roughly 1,4 kb and to contain several stretches of consecutive G residues (6). Consistent with previous observations (4, 6), we found that this 1,4 kb portion of *Airn* forms co-transcriptional R-loops after *in vitro* transcription using the bacteriophage T3 RNA Polymerase. As reported previously (6), R-loop formation on circular templates was associated with a pronounced shift in plasmid mobility on agarose gels towards the relaxed form (Figure 1A). The formation of stable and RNase H-sensitive DNA:RNA hybrids at *Airn* was confirmed by dot blots with the S9.6 antibody (15) (Figure 1B). When restriction enzymes were used to linearize the circular templates after *in vitro* transcription, R-loop formation resulted in the slower mobility of the linear template on agarose gels (Figure 1C). Strikingly, treatment with nuclease P1, which specifically digests ssDNA, completely reverted this upward shift without interfering with the stability of DNA:RNA hybrids (Figure 1C&D). These observations strongly suggest that the slower mobility induced by R-loop formation on linear *Airn* templates is caused by the extruded ssDNA and not by the presence of DNA:RNA hybrids. This is consistent with the idea that the formation of DNA:RNA hybrids at *Airn* allows the non-template strand to form secondary structures, which are responsible for the mobility shift of the linear template on agarose gels.

**Figure 1:**
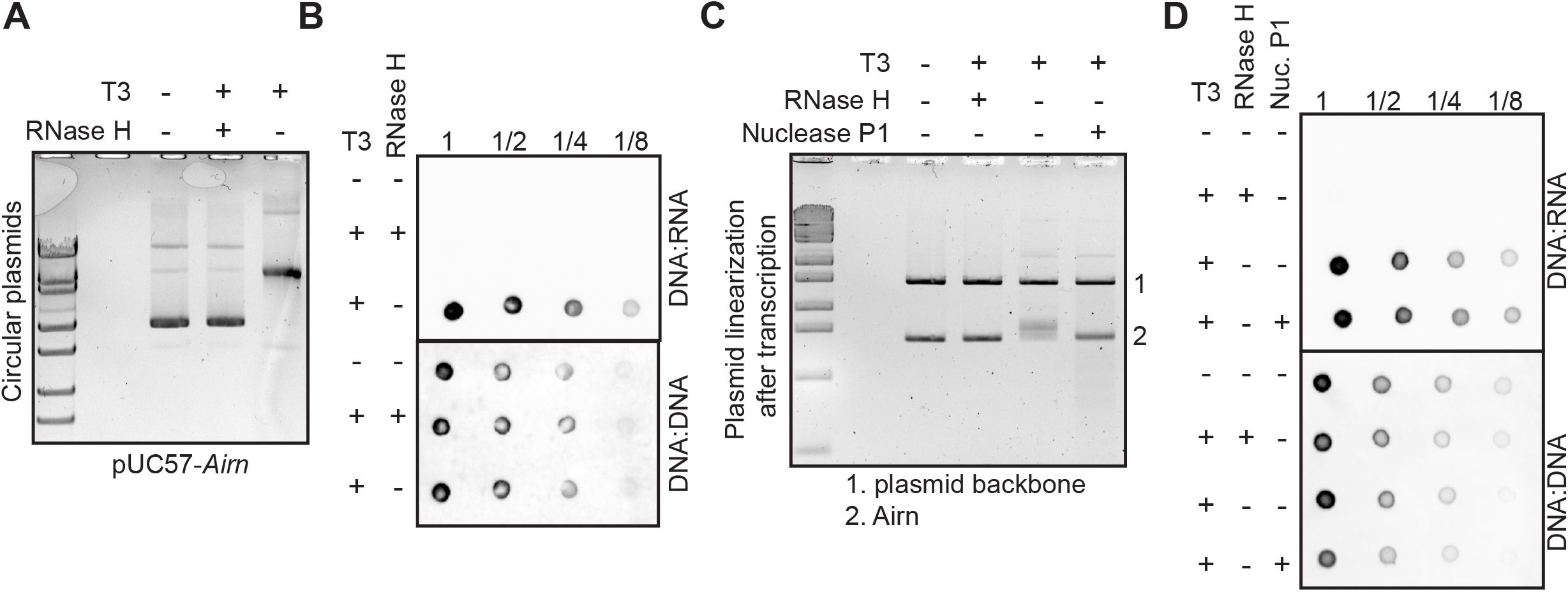
R-loop formation at *Airn* allows the folding of the non-template strand into secondary structures *in vitro*. **(AB)** The circular plasmid pUC57-*Airn* was transcribed *in vitro* and treated or not with RNase H as indicated. After purification, the DNA was run on agarose gels (A) or spotted on a membrane to perform dot blot analysis with the indicated antibodies (B). **(CD)** The circular plasmid pUC57-Airn was transcribed *in vitro* before restriction enzymes were used to separate the plasmid backbone from the *Airn* gene. After restriction digest, the reactions were treated either with RNase H or nuclease P1 as indicated. After purification, the DNA was run on an agarose gel (C) or spotted on a membrane to perform dot blot analysis with the indicated antibodies (D).

### Characterization of R-loop architecture at *Airn* using Atomic Force Microscopy

To get a better insight into the architecture of co-transcriptional R-loops formed at *Airn*, we used Atomic Force Microscopy (AFM) on mica surfaces to characterize the consequences of R-loop formation on the *Airn* template. AFM is particularly well-suited to determine with high statistics the shape and the height of individual DNA molecules at the nanometre scale (16, 17). Maximum R-loop formation was achieved by transcribing circular *Airn* templates, which were then cut using restriction enzymes to dissociate the *Airn* coding region (short fragment) from the plasmid backbone (long fragment). Both fragments were subsequently purified and imaged together on mica surfaces (Figure 2A). Strikingly, *in vitro* transcription resulted in the formation of regions of greater height and complex architecture on >80% of short fragments (Figure 2A&C). Similar objects were observed when *Airn* was transcribed from another plasmid, pFC53 (6) (Figure 2B). Regions of greater height were however rarely observed on the plasmid backbone, or on the *Airn* fragment when the DNA had not been transcribed (see below). Furthermore, their frequency was significantly reduced when transcription was followed by a treatment with RNase H to disassemble DNA:RNA hybrids (Figure 2C). Taken together, these observations demonstrate that the regions of greater height observed at *Airn* upon transcription resulted from the formation of R-loops. Thereafter, such structures are referred to as ‘R-loop objects’.

**Figure 2:**
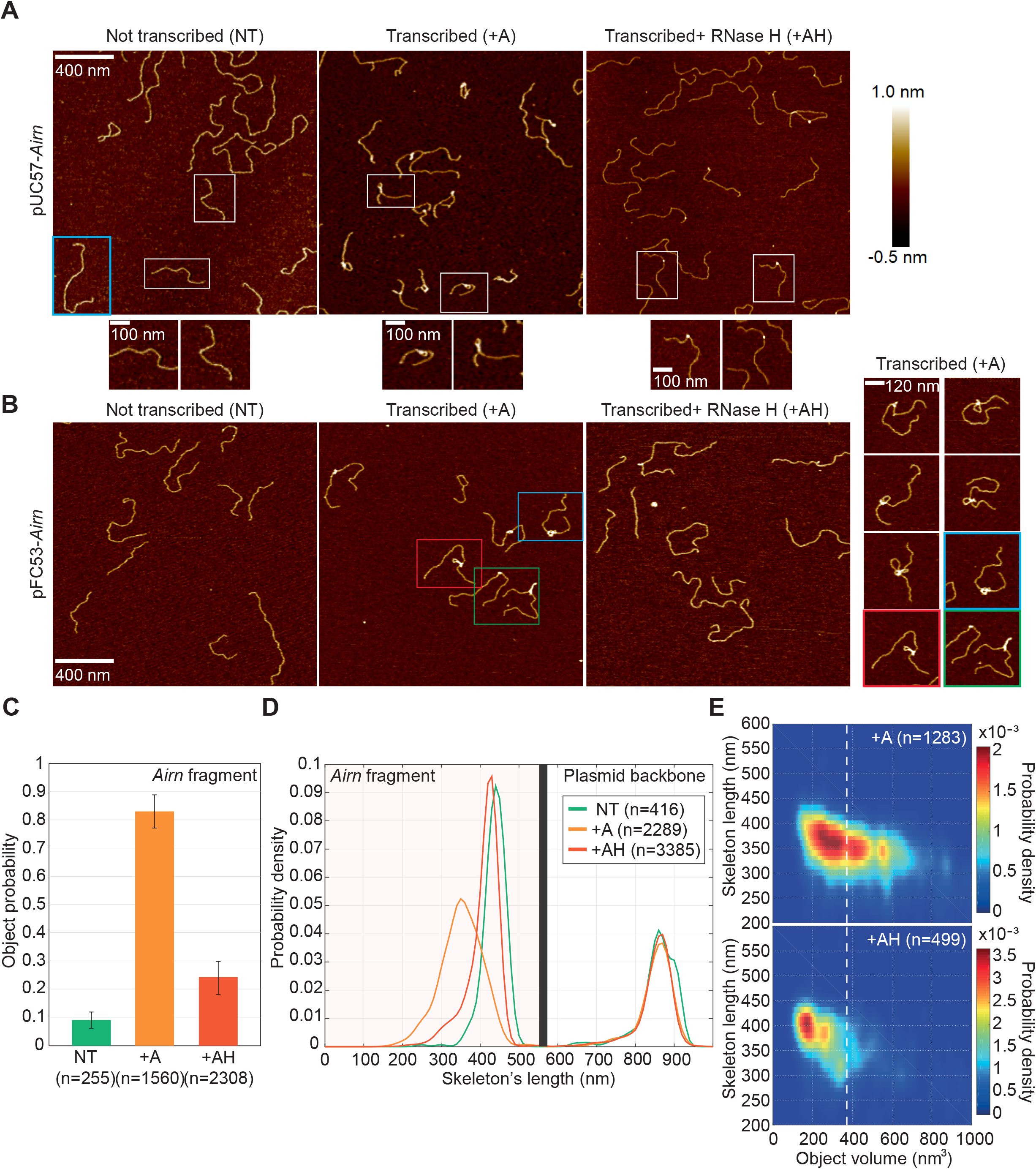
AFM analysis of the R-loop dependent secondary structures formed at *Airn*. **(A)** pUC57-Airn were processed as in Figure 1C and visualized using AFM. In pUC57-Airn, *Airn* is present on the short fragment. The blue square highlights a long DNA fragment corresponding to the plasmid backbone and white squares highly short DNA fragments corresponding to *Airn*. Magnifications of the molecules identified by a white square are shown underneath. **(B)** (left) pFC53 plasmids were processed as in Figure 1C and visualized using AFM. In pFC53, *Airn* is present on the long fragment. (right) Examples of long DNA fragments with R-loop objects. **(C)** After AFM imaging of pUC57-Airn, the probability of short molecules showing R-loop objects was established (n = 4 independent experiments; error bars represent the standard deviation). The total number of short fragment molecules counted is indicated. **(D)** Probability density of the DNA skeleton length of all the molecules analysed (both long and short fragments) in the indicated conditions (with and without objects). **(E)** 2D probability density plots showing the correlation between the skeleton length and the object’s volume in the indicated conditions.

Using a custom-built image analysis software (see methods and Figure S2), we accurately measured the skeleton length of all the DNA molecules imaged (Figure 2D). These measurements confirmed that transcription did not affect the length distribution of the plasmid backbone, consistent with our observation that R-loop objects did not form on the plasmid backbone. On the contrary, transcription resulted in the significant shortening of the *Airn* fragment. Strikingly, this shortening was largely reverted after RNase H treatment, establishing that R-loop objects assemble in a reversible and DNA:RNA hybrid-dependent manner. By making correlations between the volume of every object and the skeleton length of the corresponding molecule, we showed that R-loop objects of greater volume trigger greater reductions in skeleton length (Figure 2E), establishing that bigger objects contain more DNA. On the other hand, the few objects that remained after RNase H treatment had a significantly smaller volume (Figure 2E), consistent with the observation that they only resulted in a small reduction in skeleton length (Figure 2D). Taken together, these data show that *in vitro* transcription of *Airn* resulted in the formation of R-loop objects of varying sizes and containing varying amount of DNA.

We imaged several thousands of molecules over multiple experiments and reproducibly identified three different types of R-loop objects. We named those objects ‘blobs’, ‘spurs’ and ‘loops’ in keeping with the names used to describe the objects that formed after *in vitro* transcription of the R-loop-prone murine Sγ3 immunoglobulin switch regions (10). ‘Blobs’ represent R-loop objects aligned on the main axis of the DNA molecule, whilst ‘spurs’ come away from this axis; ‘Loops’ correspond to objects formed when a ‘blob’ sits at the base of a loop of DNA of regular height (Figure 3A). We found that ‘spurs’ were slightly more abundant than ‘blobs’ and ‘loops’ (Figure 3B) and had a greater volume (Figure 3C). Accordingly, ‘spurs’ were associated with a greater reduction in skeleton length than ‘blobs’ (Figure 3C) and therefore contained a greater amount of DNA (Figure 3D). ‘Blobs’ and ‘loops’ had similar volumes but ‘loops’ contained more DNA because they contain a loop of DNA of regular height in addition to a blob-like object. Overall, this reveals that co-transcriptional R-loops formed at the *Airn* locus consistently adopt three distinct types of three-dimensional architectures.

**Figure 3:**
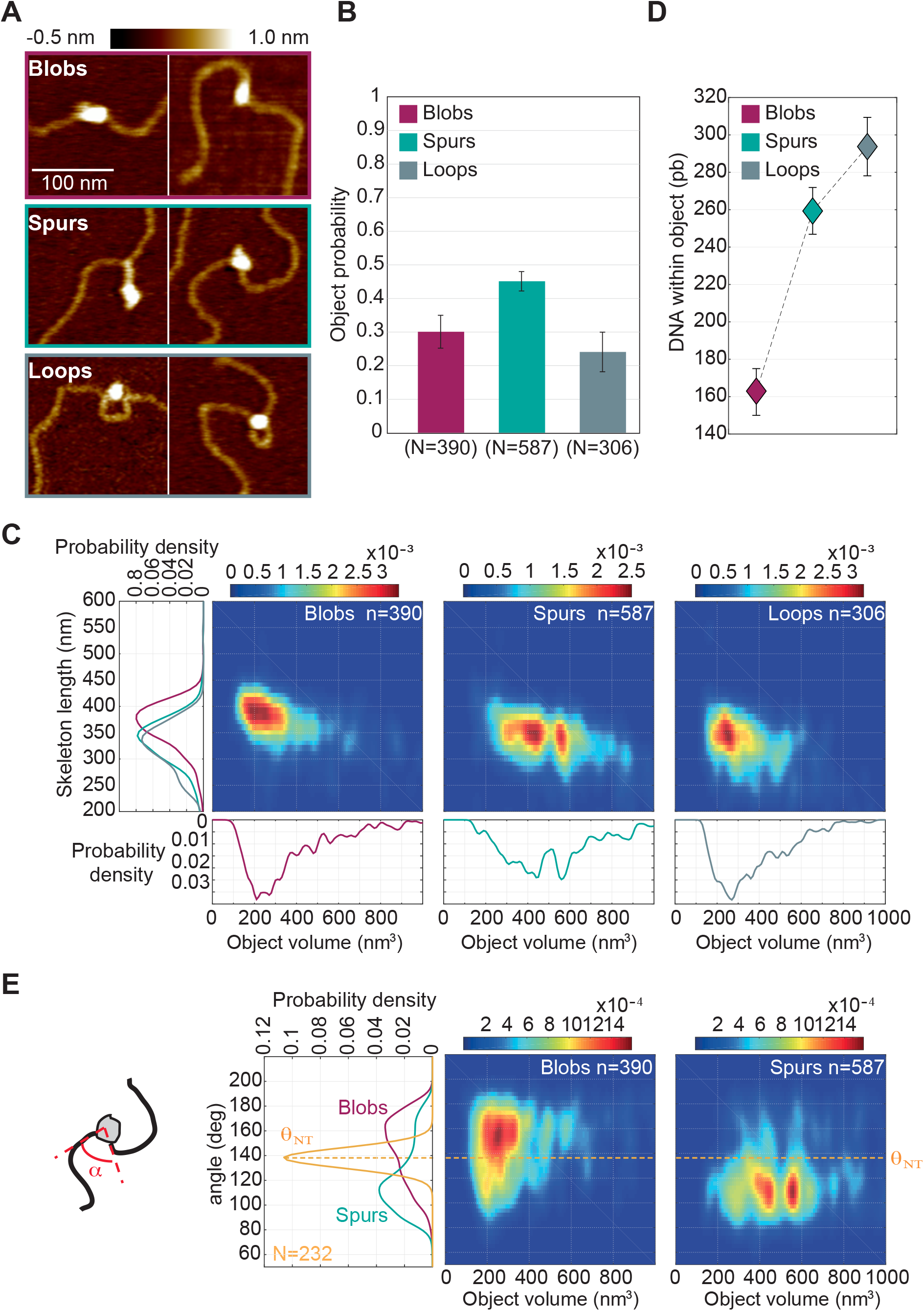
R-loop formation at *Airn* imposes mechanical constraints on the surrounding DNA. **(A)** Typical AFM images showing three different types of R-loop objects: blobs, spurs and loops. **(B)** Probability of the different types of R-loop objects (n = 4 independent experiments; error bars represent the standard deviation). **(C)** 2D probability density plots showing the correlation between the skeleton length and the object’s volume for each category of R-loop objects. The respective 1D projections of each probability density function are also shown. **(D)** Estimated amount of DNA caught in each type of R-loop objects (mean ± standard error). **(E)** 2D probability density plots showing the correlation between the angle and the object’s volume for ‘blobs’ and ‘spurs’. The angle formed between the DNA strand going into the object and the DNA strand coming out of the object was measured for ‘blobs’ (purple) and ‘spurs’ (green). The distribution of angles for molecules without objects was also established (yellow, see methods).

### R-loop objects impose local physical constraints on the surrounding DNA

We noticed that the presence of R-loop objects often introduced a marked angle in the DNA template, suggesting that R-loop objects mechanically constrain the surrounding DNA. To formalize this observation, we measured the distribution of angles formed between the incoming and the outgoing DNA strands for ‘blob’ and ‘spurs’ objects (see methods). We did not consider the ‘loop’ objects in this analysis, as it was often harder to determine the incoming and the outgoing DNA strands for this type of objects. We established that the distributions of angles for both ‘blob’ and ‘spurs’ were significantly different from the normal angle distribution determined on molecules without objects (Figure 3E), suggesting that R-loop objects kink the DNA molecule locally. Strikingly, ‘blob’ and ‘spurs’ objects introduce a different range of angles in the DNA template (respectively, 161 ±12 degrees and 110 ± 14 degrees, mean ± standard deviation). Importantly, this is irrespective of their volume (Figure 3E), which suggests that the internal organization of ‘blob’ and ‘spurs’ differs significantly. R-loop formation at *Airn* can therefore generate different types of structures, each with a distinct impact on the surrounding DNA.

To summarize, our data show that R-loop formation at *Airn* produces ssDNA-containing secondary structures that can be detected by AFM as objects of varying shape and volume. Those objects contain several hundreds of base pairs and introduce deformations in the surrounding DNA visible as kinks. As we detected several types of objects, it is likely that the extruded non-template strand of *Airn* can adopt several possible secondary structures.

### The position of R-loop objects overlaps with the position of R-loops mapped by SMRF-seq

Our observations suggested that individual transcription cycles could produce R-loops of different sequence and/or position. To accurately estimate the position of R-loop objects along the *Airn* template, we used our image analysis software to automatically measure the distance between each object and both DNA ends. Interestingly, the distance to the nearest end was similar between the different types of objects, whilst the distance to the furthest end was significantly greater for ‘blobs’ than for ‘spurs’ and ‘loops’ (Figure 4A). This observation is consistent with the idea that the starting position is conserved between objects, whilst their end position varies depending on their size.

**Figure 4:**
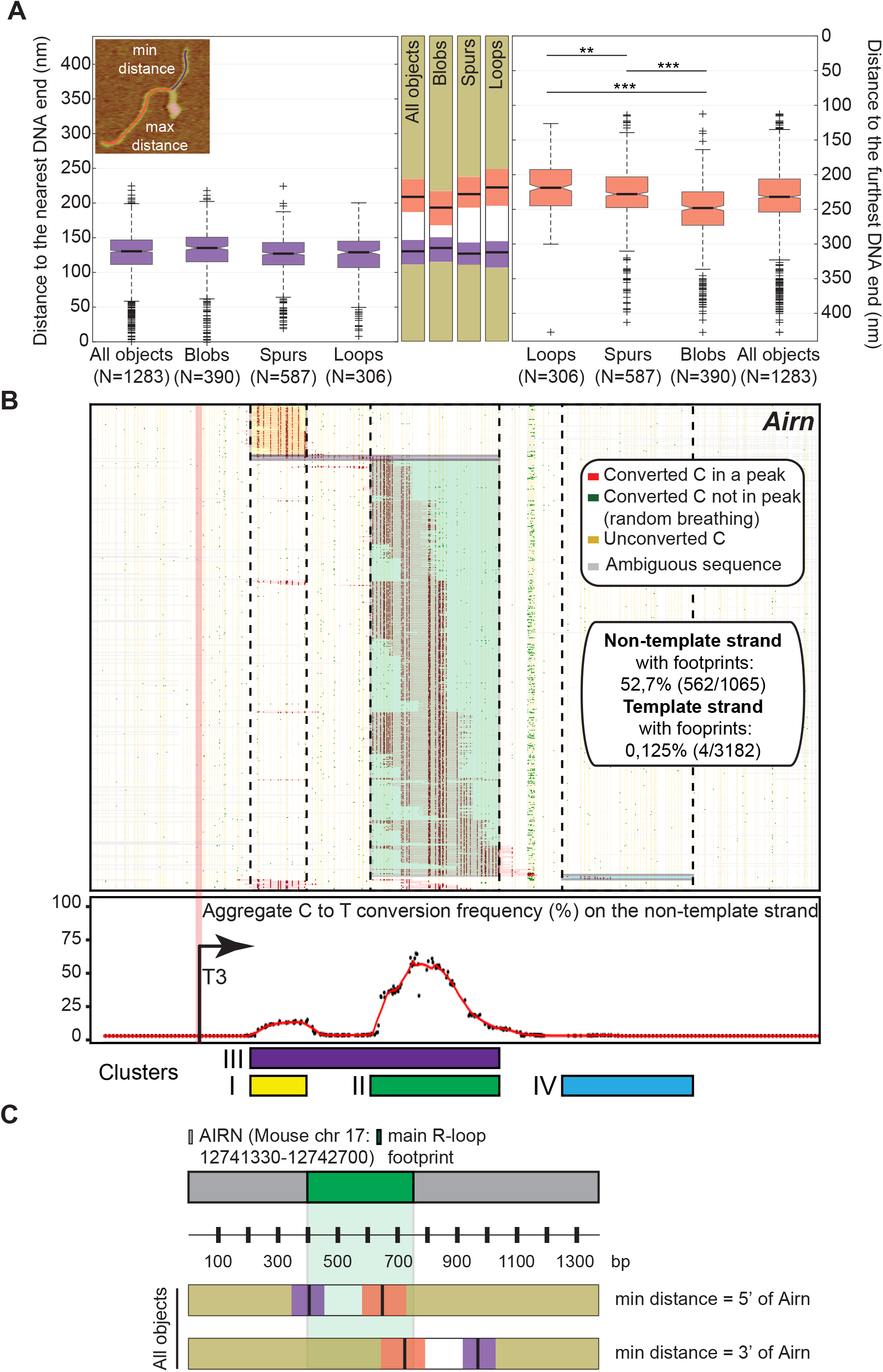
Position of R-loop objects on *Airn*. **(A)** Distance to the nearest DNA end (purple) and to the furthest DNA end (orange) measured for each R-loop object and represented as boxplots. These distances are reported schematically on the *Airn* template. The notches in the boxplots indicate the 5% uncertainty mark around the median. P-values were determined using the Wilcoxon Mann Whitney statistical test (***, p<0,001; **, p<0,01). **(B)** Footprints obtained by SMRF-seq on the non-template DNA strand. Each horizontal line corresponds to a single independent DNA molecule carrying an R-loop footprint (562 overall). The position of cytosines along the amplicon is shown by vertical orange lines. The status of each cytosine after non-denaturing bisulfite probing is color-coded as indicated in the inset; R-loop footprints are indicated by red horizontal patches. Footprints were clustered by position and clusters are highlighted using color shading. The position of the different clusters relative to the T3 promoter sequence is schematized underneath together with the aggregate C to T conversion frequency derived from R-loop peaks on the non-template DNA strand. **(C)** The position of R-loop footprints was compared to the positions of R-loop objects detected by AFM. The overlap between the two sets of measurements is best when one assumes that the nearest DNA end represents the 5’ of *Airn*.

Non-denaturing bisulfite probing can be used to map R-loops owing to the presence of bisulfite-reactive single-stranded DNA on the looped out non-template DNA strand (12). Bisulfite-mediated deamination of susceptible cytosines to uracils provides a convenient readout for singlestranded DNA that is amenable to high-throughput single molecule DNA sequencing (14). Single Molecule R-loop Footprinting (SMRF-seq) can easily be used to map R-loops generated upon *in vitro* transcription, without prior S9.6 enrichment. For this, supercoiled pUC57-*Airn* was transcribed, linearized post transcription, and R-loops mapped by virtue of the strand-specific patterns of C to T conversion induced by non-denaturing bisulfite treatment. As expected, clear strand-specific patterns of bisulfite sensitivity were observed on the displaced strand of the plasmid (Figure 4B), while the RNA-paired template DNA strand was protected from bisulfite. Based on the analysis of 562 independent R-loop-carrying molecules, R-loops were distributed principally over two clusters, with 88% of R-loop footprints mapping to cluster 2 located 422-768 bp downstream of the T3 promoter (Figure 4B). This main cluster matched well with a region of predicted high R-loop favourability (14). R-loops within cluster 2 were characterized by a discrete series of starts and stops defining overlapping molecular sub-clusters of varying sizes. The position of R-loop objects detected by AFM and the position of R-loops mapped using SMRF-seq showed striking overlap when we assumed that the nearest DNA end corresponded to the 5’ of *Airn* (Figure 4C). If we assumed instead that the nearest DNA end to the object corresponded to the 3’ of *Airn*, the overlap was poor. In addition, these data showed that the size of R-loops mapped by SMRF-seq and the size of R-loop objects detected by AFM were remarkably consistent (Figure 4C). Overall, the strong agreement between orthogonal data types, AFM mapping and chemical footprinting strongly supports the idea that R-loop objects detected by AFM correspond to genuine R-loops formed along the DNA.

### R-loop objects are resistant to the presence of lithium

Similar R-loop objects were previously reported after *in vitro* transcription of the R-loop prone murine Sγ3 immunoglobulin switch regions (10). In this case, the formation of objects at Sγ3 repeats was attributed to the assembly of the extruded ssDNA into G-quadruplexes (G4s), because their formation was sensitive to the presence of lithium ions during transcription. To evaluate the possibility that the objects we detected at *Airn* were the result of G4 formation, we carried out *in vitro* transcription in the complete absence of NaCl, which was replaced by either 40 mM KCl or 40 mM LiCl to respectively stabilize or destabilize G4, as previously described (10). The presence of LiCl had no effect on the transcription-induced shift of the DNA template on an agarose gel (Figure 5A), on the amount of DNA:RNA hybrids produced during transcription (Figure 5B) or on the abundance (Figure 5C), properties and positions of the R-loop objects detected using AFM (Figure 5D&E). Similar conclusions were reached when LiCl was added during transcription of pFC53 (Figure S3). To directly probe G4 formation at *Airn*, we used the BG4 antibody that recognizes different structural conformations of G4 (18). Dot blot assays with the BG4 antibody confirmed LiCl-sensitive G4 formation at *c-myc* and G4 formation in a plasmid containing Sγ3 immunoglobulin switch regions. Treatment with nuclease P1 to digest the G4-containing ssDNA eliminated the BG4 signal at Sγ3 immunoglobulin switch regions. Taken together, these data highlight the specificity of the BG4 antibody, as reported previously (18). However, these experiments also established that G4 formation at *Airn* was not significant and importantly not induced by transcription or sensitive to RNase H treatment (Figure 5F). We conclude that G4 formation is unlikely to explain the formation of R-loop objects detected by AFM at *Airn*.

**Figure 5:**
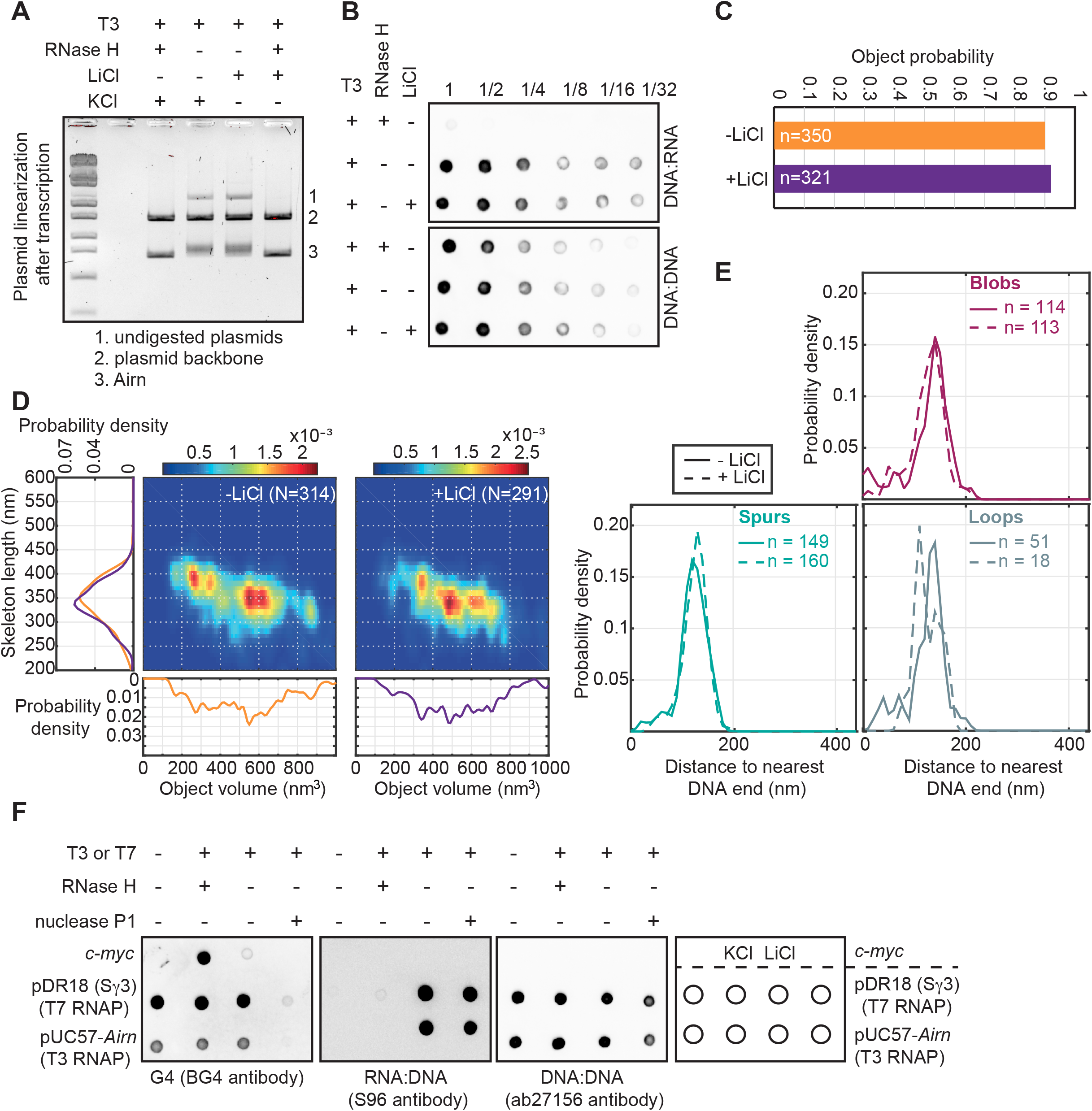
R-loop objects at *Airn* do not result from G-quadruplex formation. **(AB)** The circular plasmid pUC57-Airn was transcribed *in vitro* in the presence or not of 40 mM LiCl, before restriction enzymes were used to separate the plasmid backbone from the *Airn* template. After purification, the DNA was run on an agarose gel (A) or spotted on a membrane to perform dot blot analysis with the indicated antibodies (B). **(C)** Probability of R-loop objects. **(D)** 2D probability density plots showing the correlation between the skeleton length and the object’s volume in the indicated conditions. The respective 1D projections of the probability density function are also shown. **(E)** Distance between R-loop objects and the nearest DNA end in the indicated conditions and for the different types of objects. **(F)** Dot blot analysis of *in vitro* transcription products of pDR18-Sγ3 or pUC57-Airn with the indicated antibodies. As expected, the G4 signals disappear upon treatment with nuclease P1 and G4 formation at *c-myc* is disrupted in the presence of LiCl. See text and methods for details.

### Directed mutagenesis impacts the formation of R-loop objects without interfering with the formation of DNA:RNA hybrids

If G4 formation does not explain the formation of R-loop objects at *Airn*, we considered the possibility that they resulted from the formation of other, lithium-resistant secondary structures in the extruded ssDNA. The non-template strand of *Airn* contains a stretch of 14 G residues towards the 5’ border of the main R-loop forming region (cluster II). To test the possibility that R-loop formation would allow this G-stretch to form secondary structures by interacting with other parts of the gene through either Watson-Crick or Hoogsteen bonds, we sought to mutagenize this motif without interfering with the R-loop forming ability of *Airn*. We found that the introduction of five G to A mutations in the G stretch (Figure 6A) did not affect the amount of DNA:RNA hybrids produced after *in vitro* transcription (Figure 6B) but reduced dramatically the mobility shift of the linear *Airn* template on agarose gel (Figure 6C), in a similar way to the nuclease P1 treatment (Figure 1C). This compelling observation confirmed that the transcription-dependent and RNase H-sensitive mobility shift of linear *Airn* templates on agarose gels is not due to the formation of DNA:RNA hybrids in itself but most likely to the formation of secondary structures within the extruded ssDNA. In addition, this observation strongly suggested that the stretch of 14 G residues at the 5’ of cluster II is somehow involved in the formation of these secondary structures.

**Figure 6:**
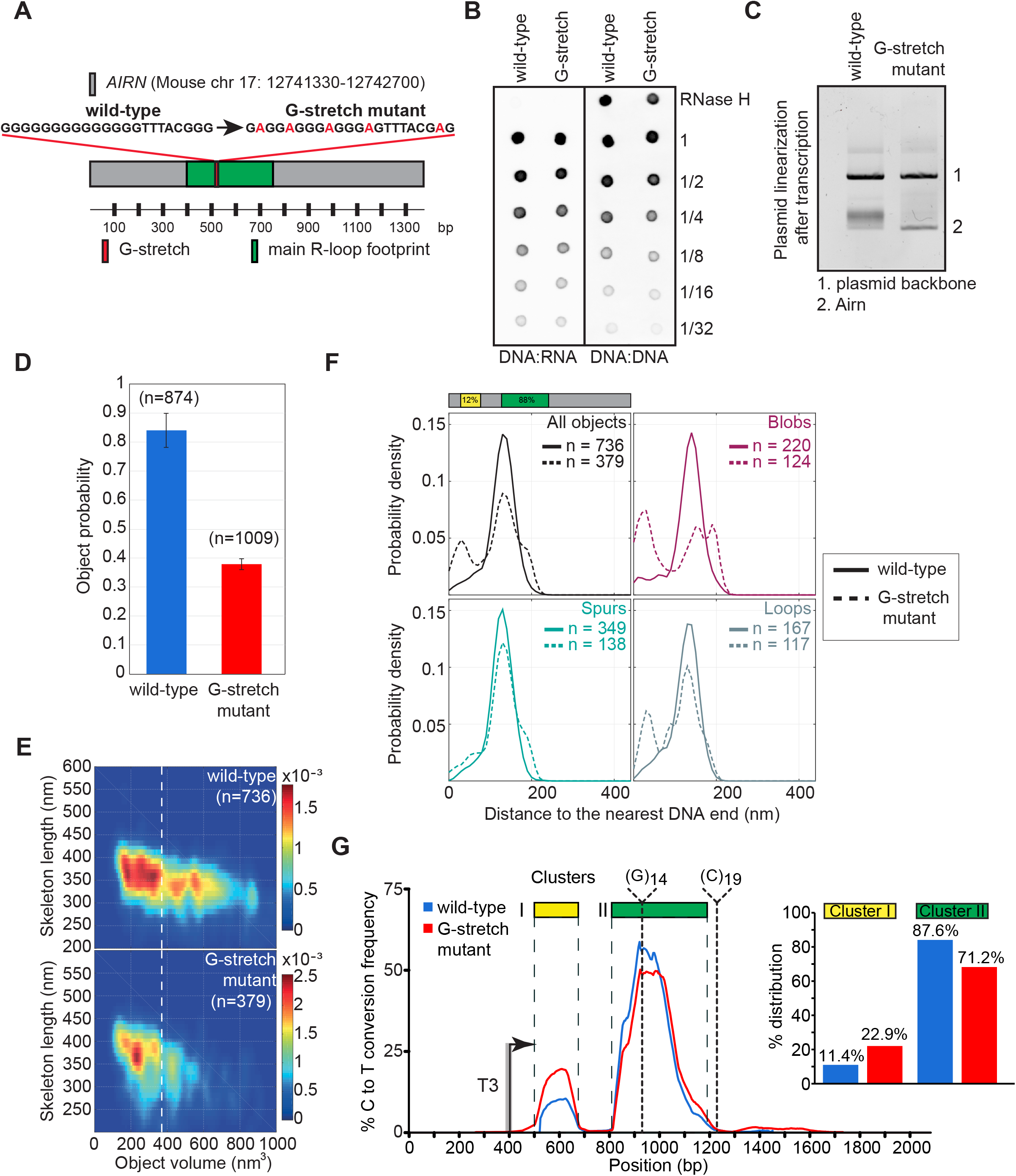
The formation of R-loop objects at *Airn* is dictated by the DNA sequence. **(A)** Schematic representation of the mutated *Airn* locus. (BC) The indicated circular plasmids were transcribed *in vitro* before restriction enzymes were used to separate the plasmid backbone from the *Airn* gene. After purification, the DNA was spotted on a membrane to perform dot blot analysis with the indicated antibodies **(B)** or run on an agarose gel (C). **(D)** Probability of R-loop objects (mean ± standard deviation, n=2). (E) 2D probability density plots showing the correlation between the skeleton length and the object’s volume in the indicated conditions. **(F)** R-loop footprinting in the G-stretch mutant. The graph displays the aggregate bisulfite conversion along the *Airn* sequence. The histograms quantify the percentages of footprints overlapping with cluster I and II in the two conditions. **(G)** Distance between R-loop objects and the nearest DNA end in the indicated conditions and for the different types of objects.

In keeping with this conclusion, the G-stretch mutations affected significantly the formation of R-loop objects detected by AFM: although the mutant template was still able to produce ‘blobs’, ‘spurs’ and ‘loops’, their frequency (Figure 6D) and their volume (Figure 6E) were strongly reduced. In addition, the position of the remaining R-loop objects was slightly altered. In particular, ‘blob’ and ‘loop’ objects were more often detected closer to the nearest DNA end in the mutant (Figure 6F), whilst the position of the remaining ‘spur’ objects was largely unaffected. This new upstream position of objects overlapped with the position of the minor cluster of R-loop footprints that was seen by SMRF-seq (cluster I, Figure 4B). In support of this observation, we confirmed that the corresponding 5’ end of *Airn* in isolation was indeed able to form R-loops and small R-loop objects (Figure S4).

Importantly, R-loop footprinting using SMRF-seq indicated that the G-stretch mutations did not dramatically affect the position of R-loops, although we detected slightly more R-loop footprints over the 5’ R-loop cluster (cluster I) in the G-stretch mutant, with a corresponding decrease over the main cluster II (Figure 6G and Figure S5). These observations again demonstrate the strong agreement between AFM mapping data and chemical footprinting data. Altogether, our results establish that *Airn* contains two R-loop forming sites, among which one overlaps the G-stretch and is much more likely to harbour stable R-loops. Mutation of the G-stretch reduced slightly the likelihood of R-loop formation at this site and increased R-loop formation upstream at the secondary 5’ site. More importantly, our data establish that to mutate just 5 residues in a 1375 nt-long template had a profound impact on the three-dimensional R-loop architecture while causing only slight alterations in R-loops themselves as measured by their frequencies, positions and lengths.

### R-loop formation at *sum3* does not produce R-loop objects

Our in-depth analysis of R-loop formation at *Airn* suggested that the formation of sequence-dependent secondary structures within the non-template strand contributed to the overall architecture and physical properties of R-loops. This suggested that R-loops could be distinguished by their architecture. To evaluate the potential diversity of R-loop architectures, we sought to identify a gene that would form comparable amount of R-loops as *Airn* upon *in vitro* transcription. Our previous mapping of R-loops at near nucleotide resolution in fission yeast (19) suggested that R-loops form at the *sum3* gene. Consistent with this, *in vitro* transcription of a circular plasmid containing *sum3* resulted in the RNase H-sensitive mobility shift of the plasmid on agarose gels towards the relaxed state (Figure 7A) and in the accumulation of DNA:RNA hybrids to levels comparable to those obtained after *in vitro* transcription of *Airn* (Figure 7B). However, when the templates were linearized, R-loop formation did not result in the upward shift of the *sum3* template, contrary to what we observed on the *Airn* template (Figure 7A, lower panel). In addition, only very few and very small R-loop objects were detected at *sum3* using AFM (Figure 7C). Consistent with this, transcription did not significantly affect the skeleton length of the *sum3* template (Figure 7D). Taken together, these observations show that the extruded ssDNA at *sum3* R-loops does not form detectable R-loop objects.

**Figure 7:**
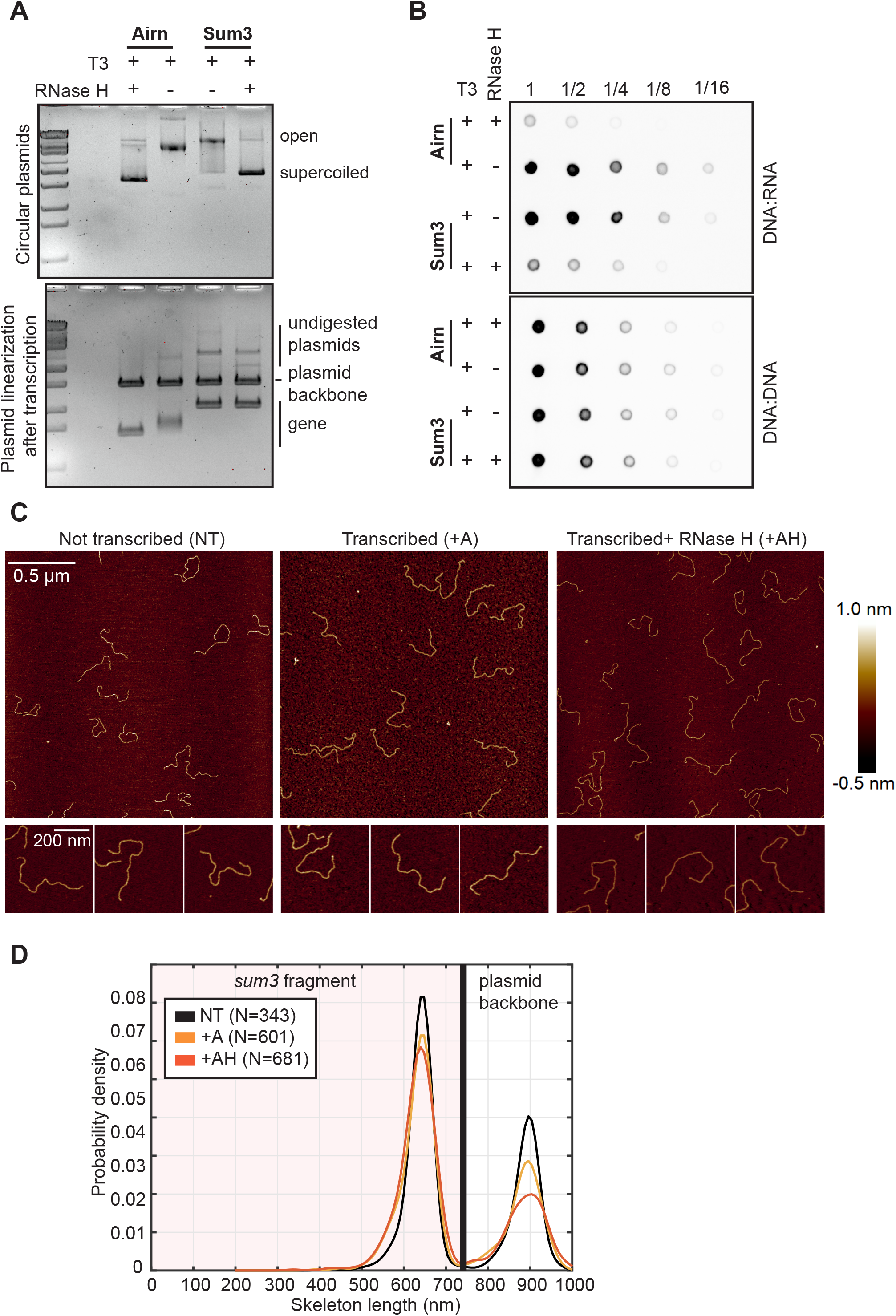
R-loop formation at *sum3* does not result in the formation of R-loop objects. **(A)** The indicated circular plasmids were transcribed *in vitro* and treated or not with RNase H and run on agarose gel before (top) or after (bottom) restriction enzymes were used to separate the plasmid backbone from the indicated gene. **(B)** The reactions from (A) were spotted on a membrane to perform dot blot analysis with the indicated antibodies. **(C)** AFM imaging of pUC57-sum3 transcribed in the indicated conditions after restriction enzymes were used to separate the plasmid backbone from *sum3. sum3* corresponds to the short fragment. **(D)** Probability density of the DNA skeleton length of all the molecules analysed in the indicated conditions.

To conclude, our results demonstrate that R-loop formation does not necessarily lead to the formation of R-loop objects detectable by AFM and that R-loops can be distinguished by their three-dimensional architecture, at least *in vitro*. In addition, we provide an easy way to identify object-forming R-loops: whilst all R-loops are able to relieve the supercoiling of a circular plasmid, we show here that only object-forming R-loops alter the mobility of linear templates on agarose gels.

## DISCUSSION

Our multi-disciplinary analysis of R-loop formation at the single molecule resolution demonstrates that the nucleotide sequence of the extruded non-template strand of R-loops determines the three-dimensional architecture of R-loops *in vitro*. Consistent with this, different R-loops adopt different conformations and to mutate the DNA sequence without interfering with R-loop formation resulted in significant alterations to the R-loop objects detected by AFM. In addition, our data are consistent with the idea that at least some R-loop architectures impose significant short-range mechanical constraints on the surrounding DNA via the introduction of kinks in the template (Figure 3E). More generally, our work suggests that R-loops possess intrinsic sequence-driven physical properties. Finally, in agreement with previous observations (10), we establish that there is significant variability in the size and shape of R-loop objects for a given gene, suggesting that individual transcription cycles produce R-loops of different sizes and architectures.

The formation of secondary structures within the extruded non-template strand of R-loops has long been hypothesized, mostly because at least some R-loops were shown to form over nucleotide sequences with structure-forming potential, as determined computationally (7, 20, 21). But to our knowledge, this possibility had never been rigorously tested experimentally. In addition, a recent study reported that only 0.4%-9% of DNA sequences with structure-forming potential actually formed detectable secondary structures *in vivo* (22). Consequently, the enrichment of structure-forming motifs in R-loops does not necessarily equate to the formation of secondary structures. We therefore surmise that only very few R-loops drive the formation of biologically relevant secondary structures and a future challenge will be to identify those R-loops that genuinely drive the formation of secondary structures *in vivo*.

What are the R-loop objects that we revealed at *Airn*? It was previously proposed that similar structures formed at the Sγ3 immunoglobulin switch regions resulted from the formation of G-quadruplex (G4) (10). In addition, it was shown recently that there is interplay between R-loop and G4 formation, at least at a subset of R-loops (23). Although the G-stretch motif that we mutated in *Airn* is predicted to be able to form G4, there are several reasons why we do not think that G4 formation underlies the structures that we detected. First, it seems unlikely that G4 formation over a 22 nt motif would be able to trigger the assembly of ~300 nt-long structures. Second, G4 formation at this motif might be predicted to form predominantly one type of structure with a given shape and a given volume, rather than the diversity of structures that we have observed. Third, our data indicate that R-loop objects at *Airn* are resistant to 40 mM LiCl, a treatment that should prevent the stabilization of G4s. Finally, no significant G4 formation was detected upon transcription of *Airn* using the G4-specific BG4 antibody. Several studies proposed that R-loop formation within a purine-rich sequence such as *Airn* could stabilize triplex H-DNA structures (H-loops) (21, 24, 25). Although this is a hypothesis that would be particularly attractive to explain the formation of the loop objects that we detected (Figure 3A), it seems inconsistent with our observations: in the G-stretch mutant, the five G to A mutations we introduced did not change the purine content of the sequence and yet they interfered significantly with the formation of R-loop objects. Further work will be required to establish exactly the type of structures that assembles at *Airn* upon R-loop formation. Whatever the exact organization of the different R-loops objects that we detected, our observations strongly suggest that R-loops might also impinge on gene expression and genome stability through the formation of other types of secondary structures than G4.

The single-stranded DNA binding protein RPA was shown previously to bind to the extruded non-template strand of R-loops and to stimulate the activity of RNase H1 against R-loops (26). It is likely that the formation of secondary structures *in vivo* will be counter-balanced by the binding of RPA to the non-template strand and it is conceivable that the half-life of an R-loop will depend directly on this dynamic equilibrium. We therefore speculate that the formation of secondary structures within the non-template strand could stabilize R-loops by limiting RNase H1 access. The fact that structure-forming R-loops could be more stable might contribute to their function and/or toxicity. When the enzymes that disassemble R-loops are defective, R-loops could have more time to form secondary structures, which could in turn alter their toxic potential and trigger genome instability. This model is consistent with the idea that only a small subset of R-loops is associated with DNA breaks (2). We speculate that structured R-loops could be more toxic for the cell and become replication impediments, as proposed recently (21). Consistent with this hypothesis, the formation of R-loops at *Airn* was shown to induce replication stress (4) and structure forming non-B DNA sequences are prominent barriers to replication fork progression (21, 27). Structured R-loops could impede directly the passage of replication forks. Alternatively, their consequences on the mechanical properties of the surrounding DNA could signal the presence of R-loops to the cell and, depending on the structures, recruit different proteins, which in turn could contribute to their toxicity. For example, it will be interesting to test whether or not structured R-loops represent better substrates for the flap endonucleases XPF and XPG that have been shown to contribute to the genotoxicity of R-loops (28). Alternatively, it was proposed that the toxicity of R-loops could be mediated at least in part by their ability to induce local alterations to the epigenome (29, 30). It is conceivable that structure-forming R-loops could have a different impact on the epigenome by recruiting different chromatin modifiers.

Finally, our data are consistent with the idea that highly structured R-loops could require more than one enzyme for complete disassembly: a DNA helicase to remove the secondary structures within the extruded ssDNA and a DNA:RNA helicase or RNase H to resolve the DNA:RNA hybrid itself (Figure S6). This could explain why we still detected small R-loop objects at *Airn* after RNase H treatment, even in the absence of detectable DNA:RNA hybrids (Figure 2). Alternatively, the complete removal of structure-forming R-loops might require specific helicases that can act against both DNA:RNA hybrid and DNA hairpins. It is unclear at this stage whether any of the helicases that have been implicated in R-loop resolution are also involved in removing secondary structures within the extruded non-template strand of R-loops. Most published evidence suggests that such helicases are more likely to target the DNA/RNA hybrid moiety of R-loops. It is however interesting to note that the FANCI/FANCD2 heterodimer was recently shown to bind to the non-template strand of G-rich R-loops *in vitro* and it was proposed that it might facilitate R-loop resolution by recruiting downstream effectors (31). It is still unclear at this stage however whether the binding of FANCI/FANCD2 heterodimers to R-loops would be influenced by secondary structures within the G-rich non-template strand. Future experiments should determine whether the proteome associated with R-loops differs whether or not their extruded non-template strand can form secondary structures.

## SUPPLEMENTARY DATA

Supplementary Data are available at NAR online.

## ACKNOWLEDGMENTS

We thank Pénélope Legros for her technical help at the beginning of this project and Dr Michael R. Lieber for reagents. We are grateful to Stella Hartono for her help with the bioinformatics analysis of SMRF-seq results. We thank Julien Gros for helpful discussions.

## REFERENCES

1. Santos-Pereira JM, Aguilera A (2015) R loops: new modulators of genome dynamics and function. Nat Rev Genet 16(10):583–597.

2. Costantino L, Koshland D (2018) Genome-wide Map of R-Loop-lnduced Damage Reveals How a Subset of R-Loops Contributes to Genomic Instability. Mol Cell 71(4):487–497.e3.

3. García-Rubio M, et al. (2018) Yra1-bound RNA-DNA hybrids cause orientation-independent transcription-replication collisions and telomere instability. Genes Dev 32(13–14):965–977.

4. Hamperl S, Bocek MJ, Saldivar JC, Swigut T, Cimprich KA (2017) Transcription-Replication Conflict Orientation Modulates R-Loop Levels and Activates Distinct DNA Damage Responses. Cell 170(4):774–786.e19.

5. Lang KS, et al. (2017) Replication-Transcription Conflicts Generate R-Loops that Orchestrate Bacterial Stress Survival and Pathogenesis. Cell 170(4):787–799.e18.

6. Ginno PA, Lott PL, Christensen HC, Korf l, Chídin F (2012) R-loop formation is a distinctive characteristic of unmethylated human CpG island promoters. Mol Cell 45(6):814–825.

7. Kuznetsov VA, Bondarenko V, Wongsurawat T, Yenamandra SP, Jenjaroenpun P (2018) Toward predictive R-loop computational biology: genome-scale prediction of R-loops reveals their association with complex promoter structures, G-quadruplexes and transcriptionally active enhancers. Nucleic Acids Res 46(15):7566–7585.

8. Roy D, Lieber MR (2009) G clustering is important for the initiation of transcription-induced R-loops in vitro, whereas high G density without clustering is sufficient thereafter. Mol Cell Biol 29(11):3124–3133.

9. Duquette ML, Handa P, Vincent JA, Taylor AF, Maizels N (2004) lntracellular transcription of G-rich DNAs induces formation of G-loops, novel structures containing G4 DNA. Genes Dev 18(13):1618–1629.

10. Neaves KJ, Huppert JL, Henderson RM, Edwardson JM (2009) Direct visualization of G-quadruplexes in DNA using atomic force microscopy. Nucleic Acids Res 37(18):6269–6275.

11. Roy D, Yu K, Lieber MR (2008) Mechanism of R-loop formation at immunoglobulin class switch sequences. Mol Cell Biol 28(1):50–60.

12. Yu K, Chedin F, Hsieh C-L, Wilson TE, Lieber MR (2003) R-loops at immunoglobulin class switch regions in the chromosomes of stimulated B cells. Nat Immunol 4(5):442–451.

13. Hänsel-Hertsch R, Spiegel J, Marsico G, Tannahill D, Balasubramanian S (2018) Genome-wide mapping of endogenous G-quadruplex DNA structures by chromatin immunoprecipitation and high-throughput sequencing. Nat Protoc 13(3):551–564.

14. Stolz R., et al. The interplay between DNA sequence and negative superhelicity drives R-loop structures. Proc Natl Acad Sci USA in press.

15. Boguslawski SJ, et al. (1986) Characterization of monoclonal antibody to DNA.RNA and its application to immunodetection of hybrids. J Immunol Methods 89(1):123–130.

16. Montel F, et al. (2011) RSC remodeling of oligo-nucleosomes: an atomic force microscopy study. Nucleic Acids Res 39(7):2571–2579.

17. Japaridze A, et al. (2017) Spatial organization of DNA sequences directs the assembly of bacterial chromatin by a nucleoid-associated protein. J Biol Chem 292(18):7607–7618.

18. Biffi G, Tannahill D, McCafferty J, Balasubramanian S (2013) Quantitative visualization of DNA G-quadruplex structures in human cells. Nat Chem 5(3):182–186.

19. Hartono SR, et al. (2018) The Affinity of the S9.6 Antibody for Double-Stranded RNAs Impacts the Accurate Mapping of R-Loops in Fission Yeast. J Mol Biol 430(3):272–284.

20. Chen L, et al. (2017) R-ChIP Using Inactive RNase H Reveals Dynamic Coupling of R-loops with Transcriptional Pausing at Gene Promoters. Mol Cell 68(4):745–757.e5.

21. Šviković S, et al. (2018) R-loop formation during S phase is restricted by PrimPol-mediated repriming. EMBO J. doi:10.15252/embj.201899793.

22. Kouzine F, et al. (2017) Permanganate/S1 Nuclease Footprinting Reveals Non-B DNA Structures with Regulatory Potential across a Mammalian Genome. Cell Syst 4(3):344–356.e7.

23. De Magis A, et al. (2019) DNA damage and genome instability by G-quadruplex ligands are mediated by R loops in human cancer cells. Proc Natl Acad Sci U S A 116(3):816–825.

24. Grabczyk E, Fishman MC (1995) A long purine-pyrimidine homopolymer acts as a transcriptional diode. J Biol Chem 270(4):1791–1797.

25. Neil AJ, Liang MU, Khristich AN, Shah KA, Mirkin SM (2018) RNA-DNA hybrids promote the expansion of Friedreich's ataxia (GAA)n repeats via break-induced replication. Nucleic Acids Res 46(7):3487–3497.

26. Nguyen HD, et al. (2017) Functions of Replication Protein A as a Sensor of R Loops and a Regulator of RNaseH1. Mol Cell 65(5):832–847.e4.

27. Shastri N, et al. (2018) Genome-wide Identification of Structure-Forming Repeats as Principal Sites of Fork Collapse upon ATR Inhibition. Mol Cell. doi:10.1016/j.molcel.2018.08.047.

28. Sollier J, et al. (2014) Transcription-coupled nucleotide excision repair factors promote R-loop-induced genome instability. Mol Cell 56(6):777–785.

29. Castellano-Pozo M, et al. (2013) R loops are linked to histone H3 S10 phosphorylation and chromatin condensation. Mol Cell 52(4):583–590.

30. García-Pichardo D, et al. (2017) Histone Mutants Separate R Loop Formation from Genome Instability Induction. Mol Cell 66(5):597–609.e5.

31. Liang Z, et al. (2019) Binding of FANCI-FANCD2 Complex to RNA and R-Loops Stimulates Robust FANCD2 Monoubiquitination. Cell Rep 26(3):564–572.e5.

