## Supplementary material for "The sequence of the extruded non-template strand determines the architecture of R-loops": Figure S

### SUPPLEMENTARY METHODS

**Image analysis.** AFM images were analysed using a home-made Matlab™ script that was developed to identify individual DNA molecules and establish whether or not they contained R-loop objects. The script could also measure the total length of the DNA skeleton, find the position of R-Loop objects on the skeleton and extract various morphological characteristics of objects (area, height, volume, angle).

*Image cleaning and molecule selection.* To discriminate between the DNA molecules and the background, we first used oversampling with the spline interpolation method then applied a bitonic filter to the image (1) before combining height and area thresholds to isolate full DNA molecules (short and long fragments) (Fig. S2A). Molecules touching the image edges were removed from the analysis.

*R-loop object selection.* R-loop objects of greater height than regular DNA molecules introduced a second peak in the height distribution plot of a typical AFM image (see Fig. S2B). To unambiguously identify R-loop objects, a series of threshold were applied. First, we used a hysteresis thresholding protocol: the height distribution was first fitted with a single Gaussian function, to obtain $h_g = m_g + 2\ s_g$, where $m_g$ and $s_g$ are respectively the mean and standard deviation of the Gaussian fit. Every pixel with a height $h > h_g$ was then removed from the height distribution and a new Gaussian function was fitted to the new height distribution (mean value m and standard deviation s). A first height threshold $h_1$ (= m + 3 s ) was applied and only pixels with $h > h_1$ were selected. This identified the tip of R-loop objects. To identify the whole object, pixels linked to those objects with a height $h > h_2$ ($h_2$ = m+ 2 s) were also considered as part of the object. Then, we only considered the objects whose minimal area was greater than 150 $nm^2$. This threshold was determined to stringently discriminate genuine R-loop objects from the height fluctuations observed on DNA molecules that are adsorbed on a flat surface. We excluded from the analysis the rare DNA molecules that contained more than one R-loop object (~1% of molecules). The type of object ('blob', 'spurs' or 'loop') was assigned manually. By comparing the results of all the AFM experiments that we carried out, we established that we cannot estimate the relative proportion of each type of objects with a better accuracy than 10% (standard deviation). For each selected R-loop object, the topographic information extracted from the image allowed us to measure precisely its height, its area and its volume.

*Length, position and angle measurements.* Each DNA molecule was skeletonized using morphological tools and its length was determined. Objects present along the molecule were excluded from the skeletonization process and we only measured the length of the free DNA arms (see Fig. S2C). To determine the position of objects, the distances to the nearest and the furthest DNA ends were measured (see Fig. S2D). We determined experimentally the conversion factor to translate nm into bp by measuring the skeleton length of the non-transcribed template (peak value at 440 nm for a 1375 bp template; conversion factor of 1 bp = 0,32 nm). Note that we could not orientate the molecules with respect to their 5' end. To determine the angle formed between the DNA strands entering and exiting an object, the DNA skeleton was first fitted with a straight line up to 5 pixel contour length distance

(~15nm) emanating from a single point on the edge of the object (maximum height value along the skeleton object). The angle a between the two lines was then measured using Al-Kashi formula: $c^2=a^2+b^2+2abcos(a)$, where a and b are the 2 DNA arm length and c the length of the third triangle side. To obtain the reference angle, the average of 5 angles measured along one molecule skeleton at random positions was considered for each molecule of DNA containing no R-Loop object (not transcribed sample).

**SUPPLEMENTARY FIGURE LEGENDS**

**Figure S1: Maps of the plasmids used in this study.**

**Figure S2: Custom-built MATLAB program to automatically identify and characterize R-loop objects. (A)** Original image and final AFM image obtained using height and area thresholding operations. **(B)** Typical probability density distribution of DNA molecule height within one AFM image containing DNA molecules with R-loop objects; AFM topographic image of a DNA molecule showing an object obtained using two different height thresholds, h1 and h2, green and pink respectively. To be selected, an object must first display a height greater than h1. Using hysteresis thresholding, the object's contour will then be extended to include all the connected pixels greater than h2. **(C)** Simplification of the DNA skeleton after removal of small junctions that result from the skeletonization process using morphological tools. **(D)** AFM topographic image of a typical DNA molecule with a R-loop object highlighting several parameters measured by the program: DNA skeleton length, distance to the nearest end (minimal distance) or to the furthest DNA end (maximal distance), angle measurement.

**Figure S3: The presence of LiCl does not impact the formation of R-loop objects when *Airn* is transcribed from the pFC53 plasmid.** The circular plasmid pFC53 containing *Airn* under the control of the T3 promoter (2) was transcribed *in vitro* in the presence or not of 40 mM LiCl, before restriction enzymes were used to separate the plasmid backbone from the *Airn* template. The resulting DNA fragments were imaged using AFM. **(A)** Probability of R-loop objects. The proportion of the different types of R-loops objects is also indicated. **(B)** 2D probability density plots showing the correlation between the skeleton length and the object's volume in the indicated conditions. **(C)** Distance between R-loop objects and the nearest/furthest DNA end in the indicated conditions (all objects).

**Figure S4: The 5' of *Airn* can form R-loops and R-loop objects. (A)** Scheme representing *Airn* and *smAirn*. **(B)** Linear fragments of *smAirn* were transcribed *in vitro* with 80 units of T3 RNA Polymerase and treated or not with RNase H. After purification, the DNA was run on an agarose gel. **(C)** AFM imaging of the transcribed templates obtained in (B). **(D)** Probability of objects. **(E)** Probability density of the skeleton length of all the molecules analysed in the indicated conditions. **(F)** 2D probability density showing the correlation between the skeleton length and the object's volume in the indicated conditions.

**Figure S5: Footprints obtained by SMRF-seq on the non-template DNA strand of the G-stretch mutant of *Airn*.** Each horizontal line corresponds to a single independent DNA molecule carrying an R-loop footprint (698 overall). The position of cytosines along the amplicon is shown by vertical orange lines. The status of each cytosine after non-denaturing bisulfite probing is color-coded as indicated in the inset; R-loop footprints are indicated by red horizontal patches. Footprints were clustered by position and clusters are highlighted using color shading. The position of the different clusters relative to the T3 promoter sequence is schematized underneath together with the aggregate C to T conversion frequency derived from R-loop peaks on the non-template DNA strand.

**Figure S6: Speculative model describing the sensitivity of the different types of R-loops to RNase H and DNA&RNA helicases.** See text for details.

**SUPPLEMENTARY REFERENCES**


1. Treece,G. (2016) The Bitonic Filter: Linear Filtering in an Edge-Preserving Morphological Framework. *IEEE Trans. Image Process. Publ. IEEE Signal Process. Soc.*, 25, 5199–5211.

2. Ginno,P.A., Lott,P.L., Christensen,H.C., Korf,I. and Chédin,F. (2012) R-loop formation is a distinctive characteristic of unmethylated human CpG island promoters. *Mol. Cell*, 45, 814–825.

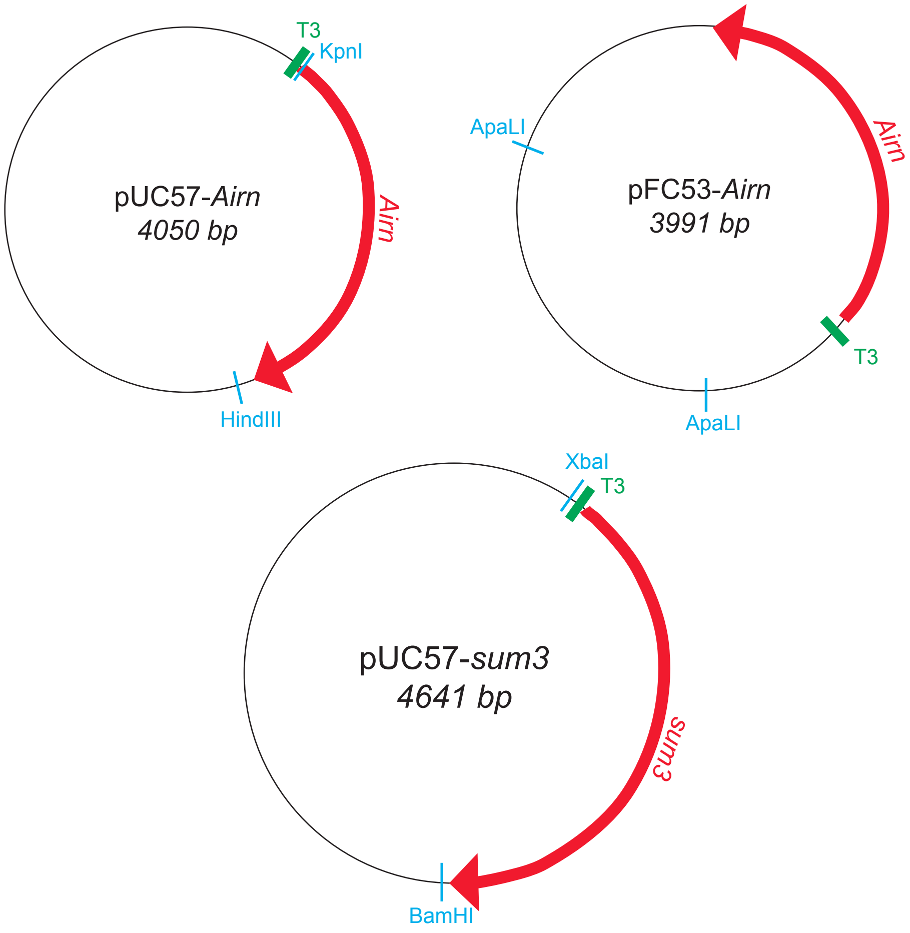
T3
KpnI
pUC57-*Airn*
*4050 bp*
*Airn*
HindIII
ApaLI
pFC53-*Airn*
*3991 bp*
*Airn*
T3
ApaLI
XbaI
T3
pUC57-*sum3*
*4641 bp*
*sum3*
BamHI

**Figure S1**

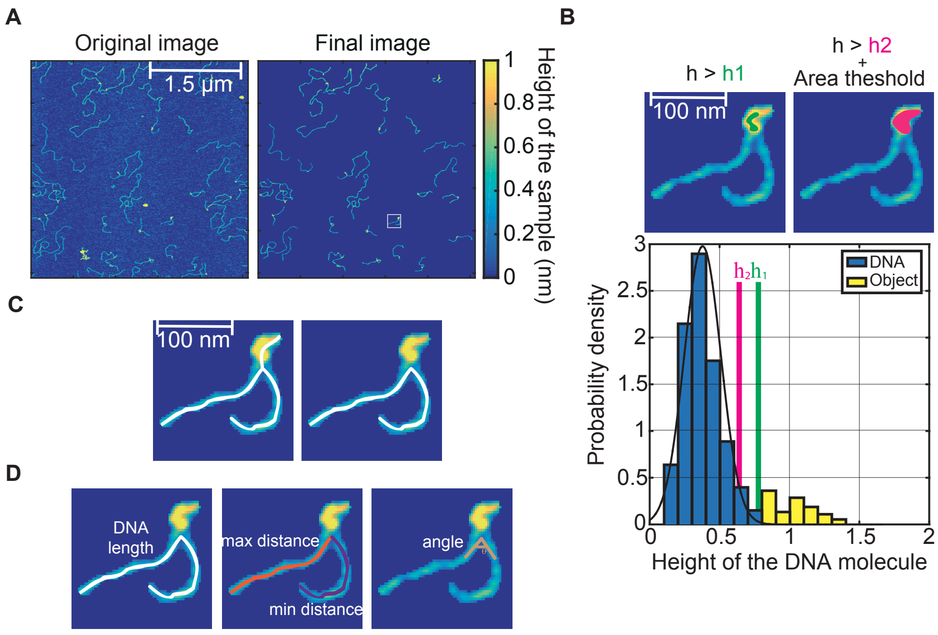
A
Original image
Final image
1.5 µm
1
0.8
0.6
0.4
0.2
0
Height of the sample (nm)
B
h > h1
h > h2
+
Area theshold
100 nm
3
2.5
2
1.5
1
0.5
0
Probability density
h2 h1
DNA
Object
0
0.5
1
1.5
2
Height of the DNA molecule
C
100 nm
D
DNA
length
max distance
min distance
angle

**Figure S2**

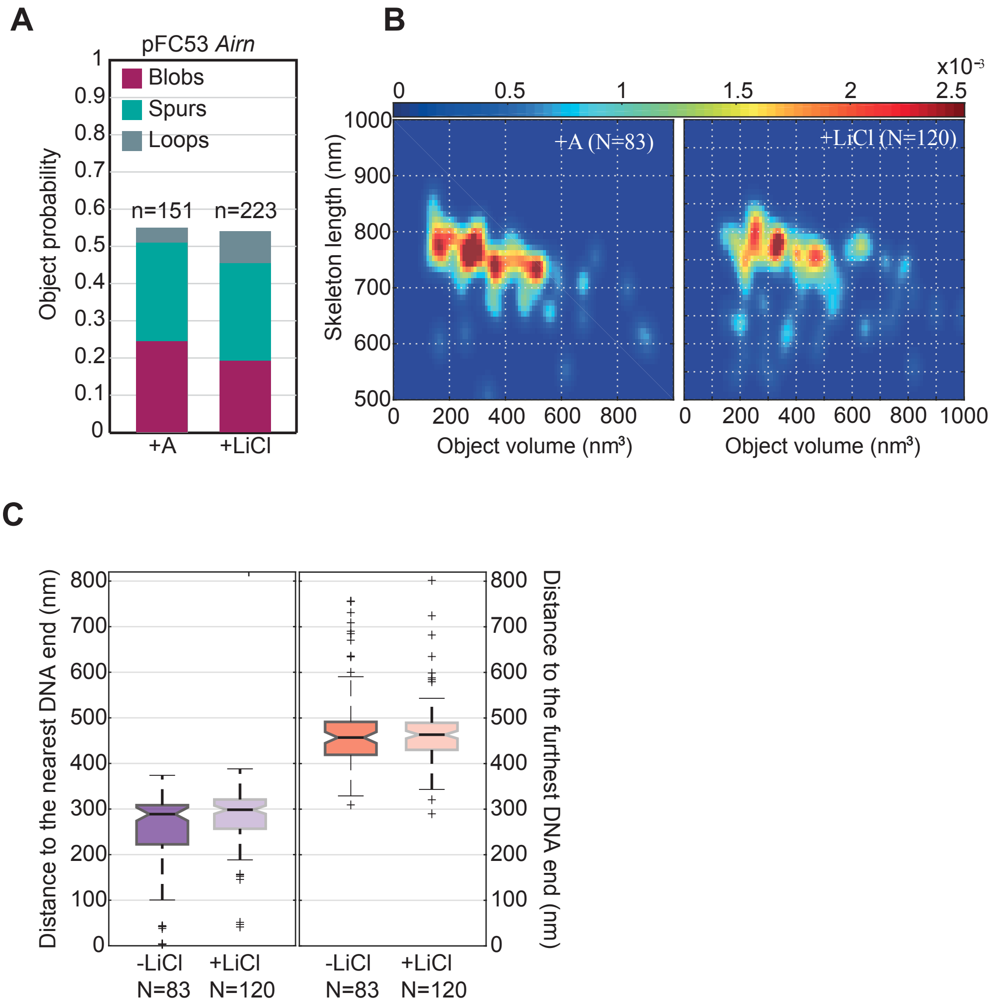
A
pFC53 Airn
Blobs
Spurs
Loops
n=151
n=223
Object probability
+A
+LiCl
B
x10-3
+A (N=83)
+LiCl (N=120)
Skeleton length (nm)
Object volume (nm³)
Object volume (nm³)
C
Distance to the nearest DNA end (nm)
Distance to the furthest DNA end (nm)
-LiCl N=83
+LiCl N=120
-LiCl N=83
+LiCl N=120

**Figure S3**

Figure S4

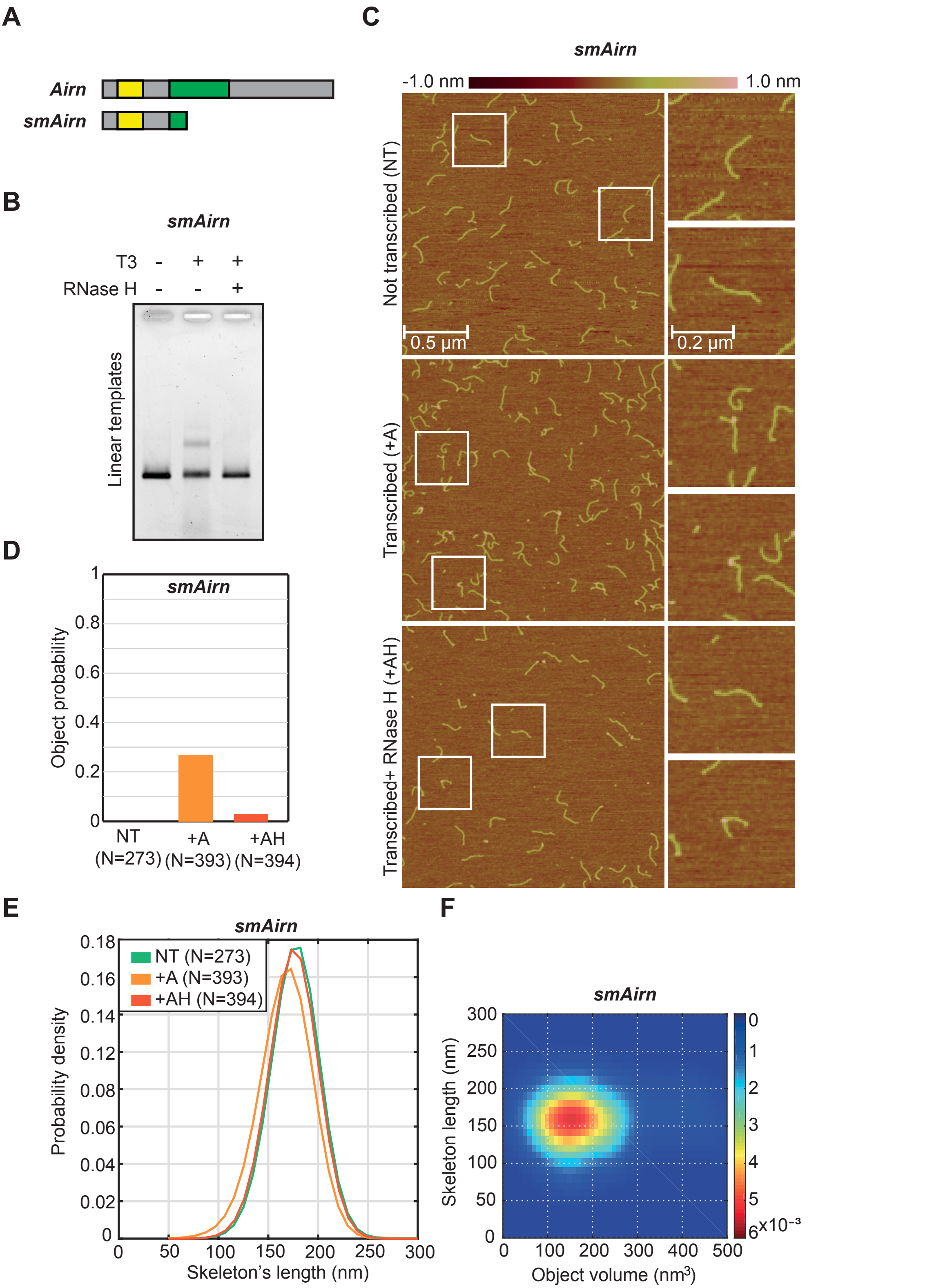
A
Airn
smAirn
B
smAirn
T3 - + +
RNase H - - +
Linear templates
C
smAirn
-1.0 nm
1.0 nm
Not transcribed (NT)
0.5 µm
0.2 µm
Transcribed (+A)
Transcribed+ RNase H (+AH)
D
smAirn
Object probability
0
0.2
0.4
0.6
0.8
1
NT (N=273)
+A (N=393)
+AH (N=394)
E
smAirn
NT (N=273)
+A (N=393)
+AH (N=394)
Probability density
0
0.02
0.04
0.06
0.08
0.12
0.14
0.16
0.18
0
50
100
150
200
250
300
Skeleton's length (nm)
F
smAirn
Skeleton length (nm)
0
50
100
150
200
250
300
Object volume (nm³)
0
100
200
300
400
500
0
1
2
3
4
5
6
x10⁻³

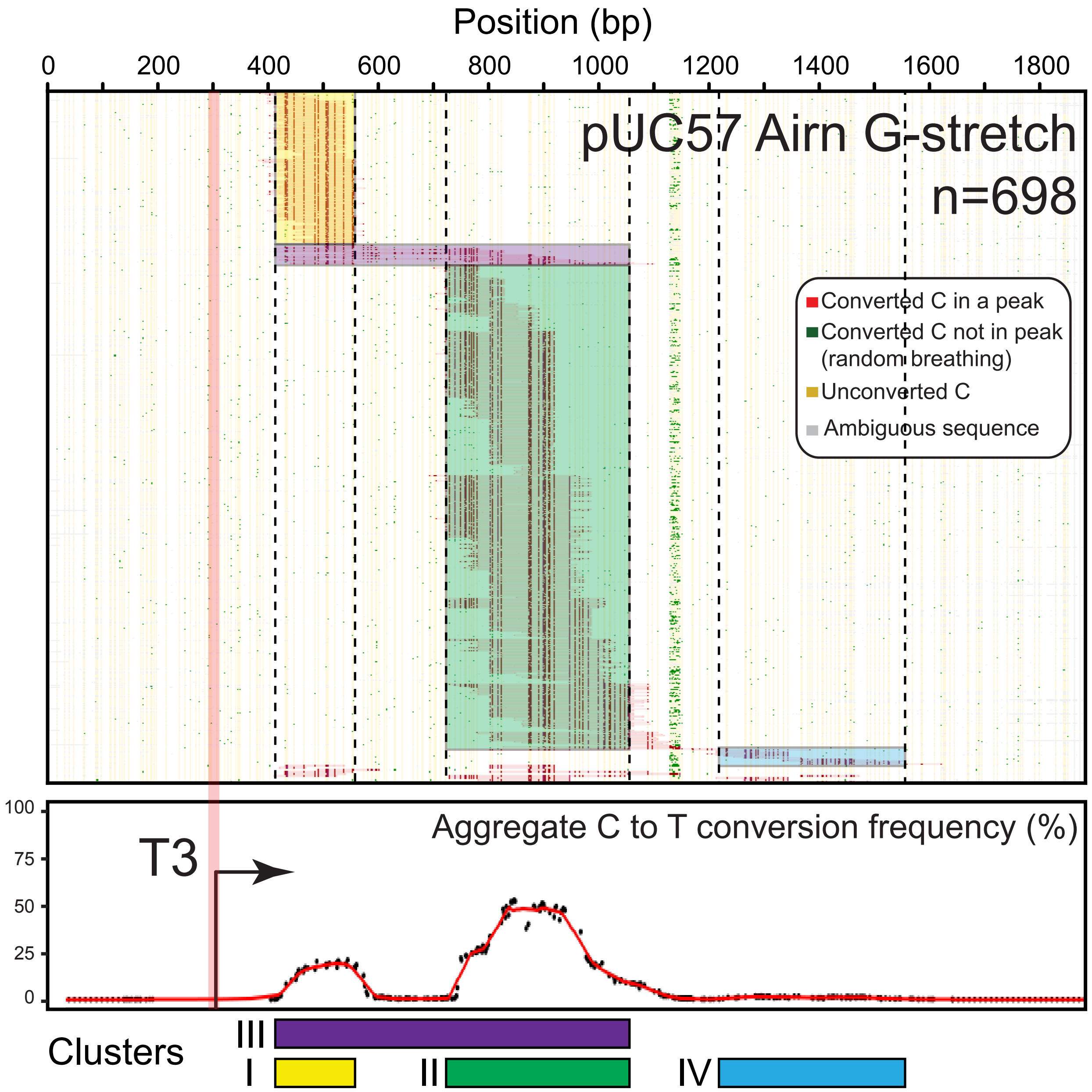

Position (bp)
0
200
400
600
800
1000
1200
1400
1600
1800
pUC57 Airn G-stretch
n=698
Converted C in a peak
Converted C not in peak
(random breathing)
Unconverted C
Ambiguous sequence
100
75
50
25
0
Aggregate C to T conversion frequency (%)
T3
Clusters
III
I
II
IV


**Figure S5**

**Figure S6**

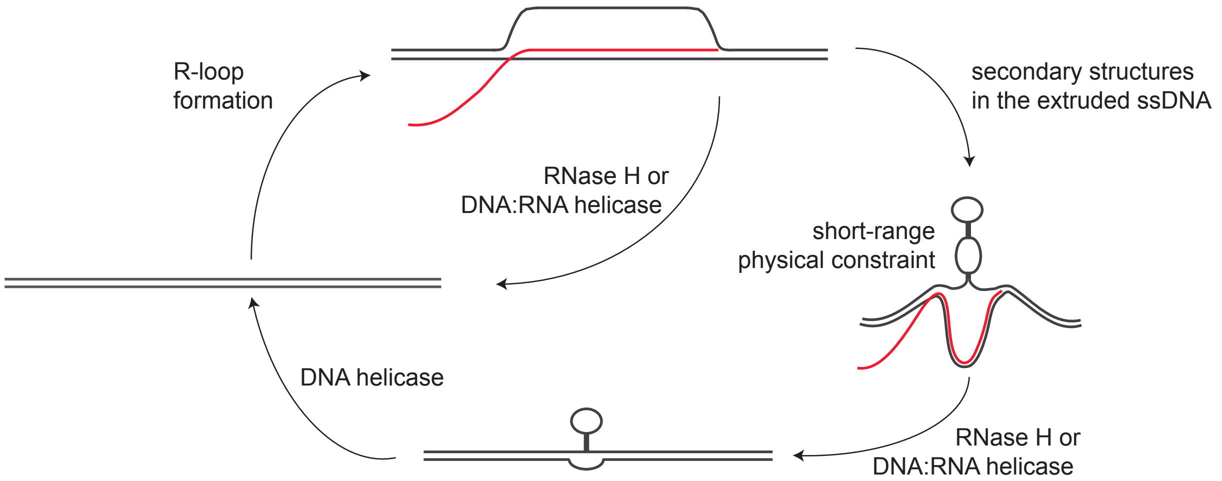

R-loop
formation
secondary structures
in the extruded ssDNA
RNase H or
DNA:RNA helicase
short-range
physical constraint
DNA helicase
RNase H or
DNA:RNA helicase
